# Distinct horizontal transfer mechanisms for type I and type V CRISPR-associated transposons

**DOI:** 10.1101/2023.03.03.531003

**Authors:** Kuang Hu, Chia-Wei Chou, Claus O. Wilke, Ilya J. Finkelstein

## Abstract

CRISPR-associated transposons (CASTs) co-opt CRISPR-Cas proteins and Tn7-family transposons for RNA-guided vertical and horizontal transmission. CASTs encode minimal CRISPR arrays but can’t acquire new spacers. Here, we show that CASTs instead co-opt defense-associated CRISPR arrays for horizontal transmission. A bioinformatic analysis shows that all CAST sub-types co-occur with defense-associated CRISPR-Cas systems. Using an *E. coli* quantitative transposition assay, we show that CASTs use CRISPR RNAs (crRNAs) from these defense systems for horizontal gene transfer. A high-resolution structure of the type I-F CAST-Cascade in complex with a type III-B crRNA reveals that Cas6 recognizes direct repeats via sequence-independent *π − π* interactions. In addition to using heterologous CRISPR arrays, type V CASTs can also transpose via a crRNA-independent unguided mechanism, even when the S15 co-factor is over-expressed. Over-expressing S15 and the trans-activating CRISPR RNA (tracrRNA) or a single guide RNA (sgRNA) reduces, but does not abrogate, off-target integration for type V CASTs. Exploiting new spacers in defense-associated CRISPR arrays explains how CASTs horizontally transfer to new hosts. More broadly, this work will guide further efforts to engineer the activity and specificity of CASTs for gene editing applications.

## Introduction

CRISPR-Cas components associate with multiple systems beyond adaptive immunity [1–3]. For example, CRISPR-associated transposons (CASTs) are an amalgam of a nuclease-inactive CRISPR effector complex and a Tn7-family transposon [4–6]. The ancestral Tn7 transposon consists of five genes, termed *tnsA-E* [6–11]. Transposition is catalyzed by *tnsA-C*, whereas *tnsD* and *tnsE* participate in target selection via two distinct mechanisms [6, 10, 11]. TnsD is a DNA-binding protein that recognizes and binds to specific sequences within the Tn7 transposon, facilitating its excision and integration into the bacterial genome. In contrast, TnsE is a structure-specific DNA-binding protein that directs Tn7 to the lagging strand during DNA replication [12]. CASTs functionally substitute both *tnsD* and *tnsE* with a CRISPR RNA (crRNA)-guided effector complex [4, 13–15]. The guiding functions of *tnsD* are substituted by “homing” spacers that target the genomic attachment site for vertical transmission [16, 17]. The mechanism of horizontal transmission, however, remains poorly understood.

CAST systems are organized into two broad categories [14–16]. Type I CASTs, which are highly related to the Tn7 transposase, use a Cascade effector complex to target transposition. By contrast, Type V CASTs evolved from a distinct Tn5053-family transposon and use Cas12k as a single RNA-guided effector protein [18, 19]. Both CAST sub-types encode CRISPR arrays that are markedly different from defense-associated CRISPR-Cas systems. First, CAST-associated CRISPR arrays are extremely short, generally fewer than three repeats [14–16]. For example, the type I-F3c system only retains a single self-targeting (“homing”) spacer, raising the question of how it can also target invading mobile DNA. In addition, type I-C CASTs do not encode any recognizable CRISPR arrays [20]. In contrast, defense-associated CRISPR arrays have tens to hundreds of repeats [21– 23]. Second, CASTs do not encode the adaptation genes *cas1* and *cas2*, suggesting that they do not update their own CRISPR arrays [22, 24–27]. Third, CASTs encode an “atypical” repeat that flanks a self-targeting spacer that is used for vertical transmission [16, 17]. These differences raise the question of how CASTs use these limited CRISPR arrays to target invading mobile elements. Alternatively, there may be one or more novel mechanisms, not previously considered, that CASTS employ during horizontal gene transfer.

Here, we show that type I and type V CASTs co-opt spacers from heterologous arrays for horizontal transmission. A bioinformatic analysis reveals that all CAST sub-types co-occur with CRISPR-Cas defense systems. Mate-out transposition assays demonstrate that type I-F, I-B, and V CASTs can use crRNAs derived from CRISPR defense systems nearly as efficiently as their own spacers. A cryo-electron microscopy structure of a type I-F TniQ-Cascade in complex with a type III-B crRNA shows that Cas6 interacts with the direct repeat (DR) of the crRNA via sequence-independent electrostatic and *π*–*π* stacking interactions. Interactions between an evolutionarily conserved Cas6 residue and a nucleotide at the apex of the DR stem-loop is essential for transposition and acts as a molecular ruler for the length of the DR stem. In agreement with this structure, we show that the DR must include a five basepair stem and a five-nucleotide loop for efficient transposition. This mechanism ensures that CASTs can mobilize into invading mobile genetic elements (MGEs) because a history of MGE infections will be updated in the active CRISPR-Cas defense locus [28]. In addition to co-opting other CRISPR arrays, type V CASTs also integrate non-specifically via a crRNA-independent copy-and-paste mechanism that requires the Cas12k effector. This process is independent of the S15 specificity factor. Our study resolves the long-standing question of how CASTs can mobilize into novel MGEs without updating their own CRISPR arrays. More broadly, we reveal design principles and potential considerations for optimizing CAST crRNAs for precision gene insertion in diverse organisms.

## Results

### CASTs co-exist with active CRISPR-Cas defense systems

We reasoned that CASTs may co-opt other CRISPR arrays that are scattered throughout the host genome for horizontal transmission. To test this hypothesis, we searched for all CRISPR arrays in the genomes of CAST-encoding organisms (Figure 1). We identified 921 genomes that encoded a CAST amongst the *∼* 1M high-quality assembled genomes in the NCBI reference sequences (RefSeq) database (Figure 1A) [20, 29]. All CASTs encoded very short or undetectable CRISPR arrays (Figure 1B). Next, we searched these CAST-encoding genomes for co-occurring CRISPR-Cas systems and orphaned CRISPR arrays. Defense systems included an active nuclease (i.e., *cas3*), adaptation genes (i.e., *cas1, cas2, cas4*), and CRISPR arrays with *∼* 10–120 spacers, suggesting active spacer acquisition (Figure 1B) [30–32]. We also observed isolated examples of “orphaned” arrays that were not adjacent to a recognizable CRISPR-Cas defense system [33–35]. Fifteen percent of genomes that encode a type I-F CAST also encode additional CRISPR-Cas systems. This statistic likely represents a lower bound on the number of co-occurring CRISPR arrays because these microbes had relatively short, highly fragmented genome assemblies in the RefSeq database (Figure S1A). All organisms with a type I-B or type V CAST encode at least one additional CRISPR array (Figure 1C) [35]. 12.5% of type I-B CASTs and 11% of type V CASTs also co-occurred with two or more additional CRISPR-Cas systems (Figure 1C). Type I-F CASTs mainly co-occurred with type III-B, I-F, I-E CRISPR defense systems. In two genomes, the type I-F CAST co-existed with a type II-A defense system (Figure 1D). By contrast, type I-B and V CASTs co-occurred with type III-B and type I-D defense systems (Figure 1D). Below, we test the hypothesis that CASTs can use protospacers from co-occurring defense-associated CRISPR arrays for horizontal transmission.

**Figure 1:**
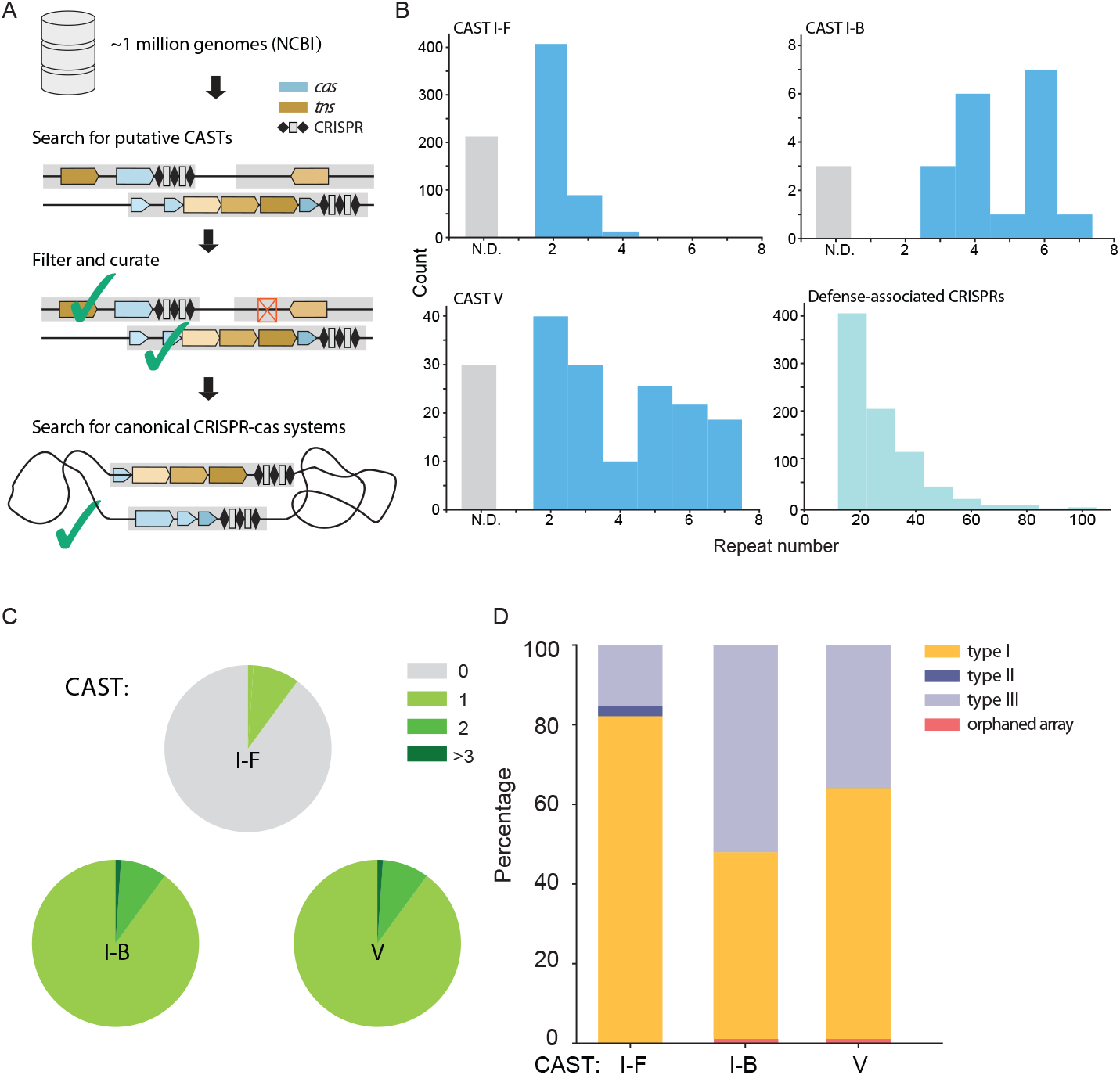
CASTs co-occur when CRISPR defense systems. (A) A bioinformatics workflow for annotating CRISPR defense systems that co-occur with CASTs in the same genome. Blue: CRISPR-associated genes; brown: transposase genes. (B) CAST CRISPR arrays are shorter than defense-associated CRISPR arrays in the same genomes. (C) CASTs frequently co-exist with one or more additional CRISPR arrays. (D) Defense-associated CRISPR-Cas sub-types that co-exist with CASTs in the NCBI microbial genome database.

### Type I-F CASTs mobilize using heterologous CRISPR arrays

To determine whether CASTs can co-opt other CRISPR arrays, we first compared the sequences and secondary structures of their direct repeats (DRs) [36]. DRs from the type I-F CAST are structurally identical to defense-associated I-F and III-B CRISPR-Cas systems, with a five nucleotide (nt) loop, five basepair (bp) stem, and five nt 3’-handle (Figure 2A). By contrast, the type I-E DR consists of a four nt loop, seven bp stem, and four nt 3’-handle. The type I-C and II-A DRs are even more divergent from the CAST I-F (Figure S1B).

**Figure 2:**
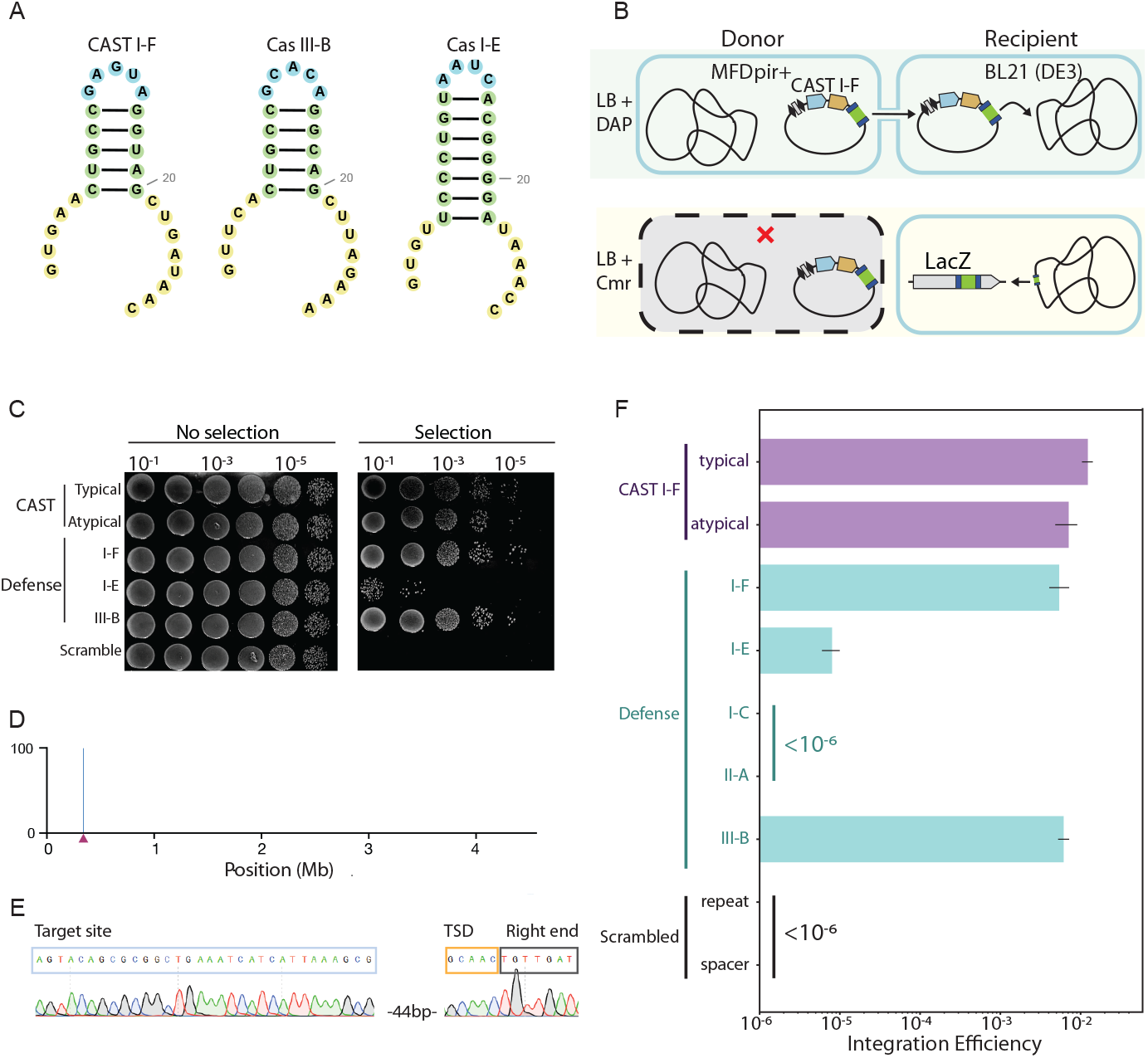
Type I-F CASTs co-opt defense-associated CRISPR arrays. (A) The predicted structures of direct repeats (DRs) from a type I-F CAST and co-occurring defense CRISPR-Cas systems. Blue: 4-5 nt loop; green: 5-7 bp stem; yellow: 5’- and 3’-handles. (B) Schematic of a quantitative conjugation-based mate-out transposition assay. A plasmid harboring the CAST, along with the cargo antibiotic resistance (green), and a minimal CRISPR array is conjugated into the recipient strain. Guided transposition into *lacZ* is scored as white, chloramphenicol-resistant clones. The donor strain is removed via counter-selection with diaminopimelic acid (DAP). (C) Direct repeats from the defense associated CRISPR arrays support transposition, but a scrambled direct repeat does not. (D) Colony-resolved long-read sequencing (E) and Sanger sequencing (F) confirms cut-and-paste transposition into *lacZ* (triangle in E). Target site duplication (TSD) is also visible in this data. (F) Quantification of transposition from the native CAST array and cooccurring defense systems. Error bars are the standard deviation across three biological replicates. Scrambling either the repeat or spacer suppressed transposition below our detection limit of *<* 10^6^ cfus.

We developed a conjugation-based chromosomal transposition assay to determine whether CASTs can exploit these heterologous CRISPR arrays (Figure 2B) [17, 37]. In this assay, the CAST genes, a CRISPR array, and a chloramphenicol (Cm) resistance marker surrounded by left and right inverted repeats are assembled into a conditionally replicative R6K plasmid that only replicates in *pir*+strains [38–40]. The *pir*+;donor also includes a chromosomally integrated RP4 conjugation system [40, 41]. Donor cells are auxotrophic for diaminopimelic acid (DAP), allowing for counter-selection on DAP-plates following conjugation with a recipient strain [39, 42]. The BL21(DE3) recipient cells support CAST expression and transposition [14, 15]. Conjugative transfer of the R6K plasmid into the recipient cells and subsequent transposition of the CAST cargo into the host genome (targeting *lacZ*) results in chloramphenicol-resistant, Δ*lacZ* recipient cells. The R6K plasmid is lost shortly after conjugation in the recipient cells (*pir-*) and the donor cells are also removed due the absence of DAP [43]. Genomic transposition efficiency can be scored quantitatively via the ratio of recipient colonies on standard (DAP-) agar plates vs. CmR cells. Targeting *lacZ* results in white colonies on Cm/X-gal plates; integration outside *lacZ* produces blue colonies on the same plates [44, 45]. Finally, we also scored the insertion accuracy via both Sanger- and whole-genome long-read sequencing.

We first tested this assay with the native and atypical direct repeats from the well-characterized *V. cholerae* HE-45 Type I-F3a system (Figure 2C) [14]. This CAST encodes an atypical direct repeat and a homing spacer for site-specific integration into the host’s genome. We removed the homing spacer to avoid spurious transposition events [17]. Transposition efficiency was scored using a *lacZ* -targeting spacer [14]. A scrambled spacer or a scrambled direct repeat served as negative controls. The transposition efficiency was 1.4 ± 0.2% of all viable recipient cells. This was suppressed below the limit of detection (< 10^*−*6^ cfus) when either the spacer or the repeat were scrambled. All chloramphenicol-resistant colonies (*n* = 395 across three biological replicates) were white on X-gal plates, suggesting transposition into *lacZ* (Figure S1C). Whole-genome long-read sequencing indicated a single transposition event at the expected target size (Figure 2D). Sanger sequencing of the insertion junctions from 32 colonies showed that the cargo inserted *∼* 42–46 bp downstream of the end of target site (Figures 2E, S2A). Integration occurred in the forward direction in 91% of all cases and in the reverse direction in the remaining 9%. An atypical direct repeat supported a nearly identical transposition efficiency and insertion orientation (Figure S2). The atypical direct repeat maintains the same overall stem-loop structure but has 12 nucleotide substitutions relative to the typical direct repeat [17]. Because the typical and atypical direct repeats maintained a high transposition rate, we conclude that the CAST effector complex can tolerate DRs with divergent RNA sequences.

Next, we tested whether this CAST can use DRs from co-occurring CRISPR defense systems (Figures 2C, F) [14]. For this assay, the native CAST array targeted *lacZ* but encoded the DR from defense-associated CRISPR-Cas systems. All other protein and cargo components remained unchanged. Surprisingly, type I-F and III-B DRs supported transposition efficiencies that were comparable to those from the native CAST, despite differing in the RNA sequence (Figure S1). CRISPR RNAs with type I-E DRs transposed ∼ 10^3^-fold less efficiently than the native CAST crRNAs (Figure 2F). In all cases, > 99% of the resulting colonies were white on X-gal plates, indicating targeted transposition into *lacZ* (Figure S1C). Integration occurred ∼ 43–45 bp from the 3’ end of the target, with 90% of all events in the forward orientation (Figures 2E. S2B). Long-read sequencing showed that a single copy of the cargo was inserted into *lacZ* (Figure 2D) [14]. By contrast, type I-C and II-A direct repeats did not support any transposition activity (< 10^*−*6^ cfus). The structures of these DRs differ substantially from the I-F DR, indicating that the DR stem loop is a major determinant of transposition.

### Cas6 stabilizes direct repeats via sequence-independent electrostatic interactions

To investigate the molecular basis for how CASTs exploit heterologous CRISPR arrays, we used cryo-electron microscopy to solve the structure of the *V. cholerae* HE-45 Cascade co-purified with a type III-B crRNA (Figure 3). The crRNA contained a native direct repeat from the type III-B system and a 32 bp spacer. The density for Cascade and the crRNA was refined with a prior model (PDB: 6PIG; Figure S3) [46–49]. The overall structure was quite similar to the prior model (*C*_*A*_ *− RMSD* = 0.83 Å), indicating the native DR from the type III-B can assemble a functional Cascade.

**Figure 3:**
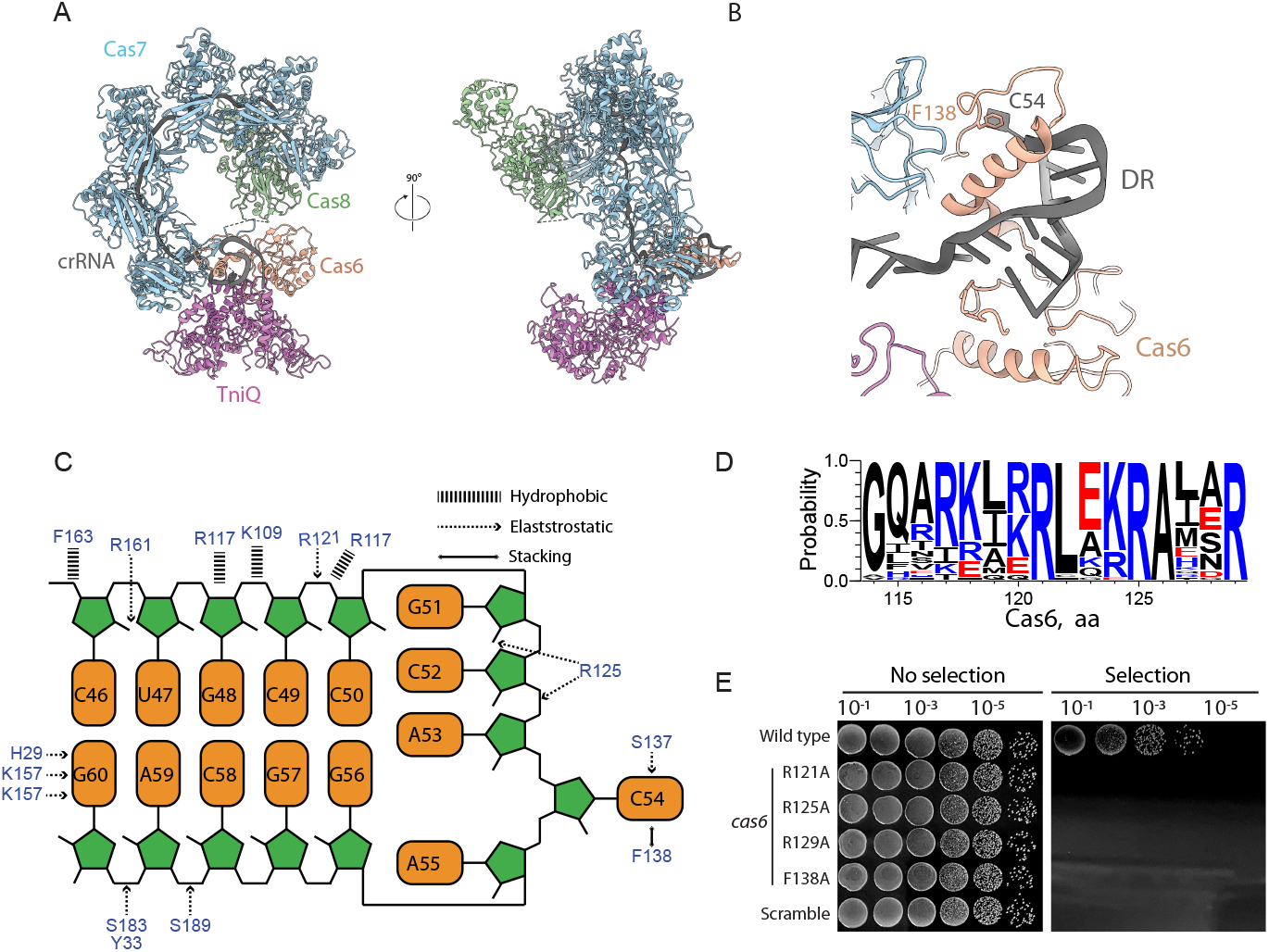
Cas6 recognizes the crRNA via sequence-independent interactions with the DR. (A) Structural overview of a type I-F TniQ-Cascade purified with a type III-B crRNA. (B) Close-up view of the direct repeat (gray) and its interactions with Cas6 (salmon). (C) Schematic of the hydrophobic and electrostatic interactions between key Cas6 residues and the direct repeat from the type III-B crRNA. (D) Multiple sequence alignment across all CAST I-F *cas6* genes reveals conserved residues in the arginine-rich helix. (E) Transposition requires Cas6 residues R121, R125, R129, and F138 to coordinate the direct repeat.

The type III-B direct repeat engages Cas6 via sequence-independent interactions with the ribose phosphate backbone (Figures 3B–C). The guanidine (G54) at the apex of the stem-loop is flipped out of the plane and enters in a long-range *π*–*π* interaction with Cas6(F138). A helix with three arginines (R117, R121, R125) also forms a strong positive pocket to stabilize the crRNA handle. A multiple sequence analysis of I-F Cas6 proteins indicates that these electrostatic interactions are conserved across the entire CAST sub-family (Figure 3D). Thus, Cascade engages diverse direct repeats via crRNA-sequence independent mechanism.

We tested the functional significance of the conserved Cas6 residues using the transposition assay described above (Figure 3D). Mutating any of the arginines to an alanine suppressed transposition below our detection range (< 10^*−*6^ cfus). Similarly, Cas6(F138A) reduces transposition > 10^4^-fold, indicating that the *π*–*π* interaction is also necessary for stably engaging the DR (Figure 3C). We conclude that Cas6 stabilizes diverse DRs via RNA sequence-independent electrostatic and *π*–*π* stacking interactions.

### The direct repeat tunes transposition efficiency

The reduced transposition efficiency with type I-E DRs indicates additional constraints on the CAST crRNA. To test these constraints, we systematically varied the DR sequence and/or structure and assayed the resulting transposition efficiency (Figure 4A). We first scrambled the DR nucleotide sequence but retained the 5 bp stem, the 5 nt loop, the 5 bp 5’ handle, and the 8 bp 3’ handle of the type I-F CAST. Surprisingly, this crRNA maintained wild type transposition efficiency (Figure 4B). By contrast, scrambling the stem-loop entirely abolished transposition. These results confirm that Cas6-DR contacts are sequence independent but require a structured DR to maintain activity.

**Figure 4:**
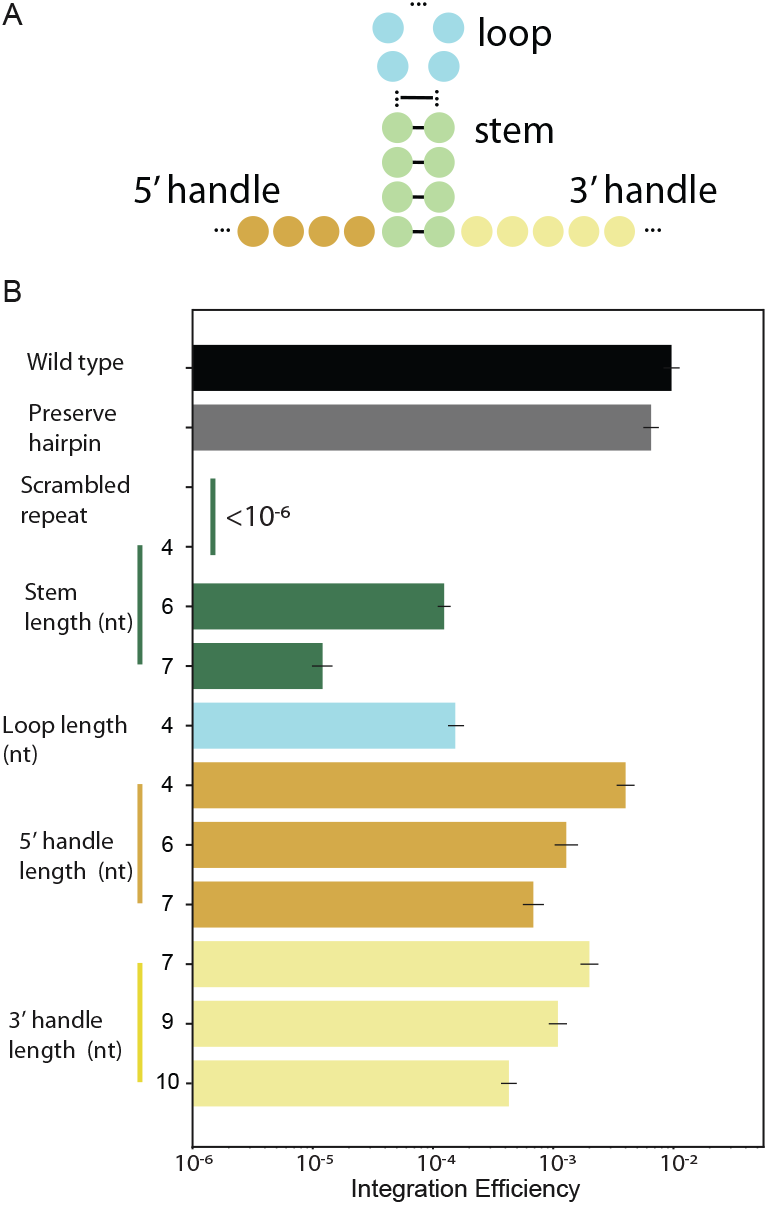
The crRNA stem-loop length regulates transposition activity. (A) We tested changes in the direct repeat sequence, stem (green), loop (blue), and handle lengths (5’-orange, 3’-yellow). (B) The effect of each feature on the integration efficiency. Black: DR from the native CAST CRISPR array; gray: a sequence-scrambled DR that preserved the wild type stem loop structure; other colors correspond to the schematic in (A).

Next, we systematically varied the length of the stem, loop, and the 5’ and 3’ handles to determine the key determinants of efficient transposition. Starting with the CAST I-F DR, changing the stem length by even a single basepair reduced transposition efficiency up to five-fold. Increasing the length of the stem from five to seven basepairs (as in the type I-E DR) decreased transposition efficiency 500-fold as compared with the type CAST I-F DR (Figure S4A). Decreasing the loop by one nucleotide also reduced transposition efficiency 100-fold (Figure 4B). Changing the length of the 5’ and 3’ handles modestly reduced transposition efficiency. Consistent with these findings, shortening the type I-E DR stem from seven to five basepairs significantly increased transposition. Adding one nucleotide from the loop to five nucleotides also improved transposition 500-fold relative to the type I-E DR (Figure S4B). These results underscore that the DR structure is the key determinant for assembling a TniQ-Cascade effector complex. The stem must be five basepairs, whereas the loop can tolerate one nucleotide changes from the five-nucleotide native sequence. The structural basis for both effects likely arises from the base stacking interaction with Cas6.

### Type I-B CASTs co-opt co-occurring CRISPR arrays for horizontal transfer

All type I-B CASTs co-occur with type I-A, I-B, I-D, or III-B defense systems or with orphaned CRISPR arrays (Figure 1C). To test whether type I-B CASTs can use these CRISPR arrays, we adapted the mate-out transposition assay to the *Anabaena variabilis* ATCC 29413 type I-B CAST (Figure S5A) [16]. We first tested transposition with the native CAST DR. Transposition efficiency with the native CRISPR sequence was ∼ 10^3^-fold lower than the type I-F CAST (Figure S5B). This may be due to poor expression in *E. coli* since we did not optimize codon usage, promoters, or translation efficiency. Most of the chloramphenicol-resistant colonies where white (∼ 90%, *n* = 152), indicating that *lacZ* was disrupted. Sanger sequencing across the insertion junctions confirmed on-target integration ∼ 44–48 bp away from the target site (Figure S5E). Scrambling the crRNA without preserving the DR structure ablated all transposition activity (Figure S5B). These results indicate that type I-B CASTs are active in the mate-out transposition assay.

Next, we tested whether the type I-B CAST can use DRs from a co-occurring CRISPR defense system. Surprisingly, the predicted structure of the co-occurring CRISPR defense system is divergent from that of the CAST (Figure S5B). The orphaned DR supported transposition, albeit ∼ 20-fold lower than the native DR (Figure S5C). We did not detect integration for any other DRs (< 10^*−*6^ cfus). We conclude that type I-B and I-F CASTs can both co-opt heterologous CRISPR arrays, so long as the crRNA DRs can be structurally accommodated within the Cascade effector complex.

### Type V CASTs transpose via crRNA-dependent and independent mechanisms

All type V CASTs co-exist with either type I or III defense-associated CRISPR systems (Figure 1). Therefore, we assayed whether the *S. hofmannii* (Sh) type V CAST can use spacers from these CRISPR arrays for horizontal transfer [15]. As before, we removed the CAST’s homing spacer and targeted the native tracr-crRNA, or a single guide RNA (sgRNA) to *lacZ*. Recent structural studies identified the small ribosomal protein S15 as a core CAST subunit that increases the on-target transposition rate [49, 50]. We therefore also tested whether the *E. coli* or *S. hofmannii* S15 aides transposition in our assay [49, 50]. Transposition efficiency was scored by measuring the colony forming units on chloramphenicol-resistant plates. On-/off-target events were confirmed via Sanger and long-read whole genome sequencing.

Transposition remained high with the native crRNA, the sgRNA, or crRNAs with type I-D direct repeats (Figure 5A). In addition, the ShCAST co-occurs with two orphaned CRISPR arrays, which also support transposition (Figure 5A). Ablation of *cas12k* dropped transposition below our detection limit (< 10^*−*6^ cfus). Surprisingly, deleting the entire crRNA (ΔcrRNA) did not diminish this activity, indicating that unguided transposition is robust with apo-Cas12k. Expressing either *E. coli* or *S. hofmannii* S15 (EcS15 or ShS15) reduced transposition ∼50–250 fold, respectively. When S15 was not over-expressed, off-target integration dominated all insertion events (Figure 5B). Over-expressing S15 in the recipient cells increased on-target integration up ∼60% of all events. Integration with the defense-associated type I-D DR was indistinguishable from the native crRNA (Figures 5C; S6). CASTs assembled with a sgRNA transposed at the target site most frequently, especially in recipient cells over-expressing ShS15. Transposition was > 95% in the L/R orientation (*n* = 613 insertion events). The cargo DNA inserted via a co-integration mechanism ∼ 29–34 bp away from the end of the *lacZ* target site (Figure 5C-E). Taken together, we conclude that ShCAST supports robust guided transposition with heterologous defense-associated CRISPR arrays.

**Figure 5:**
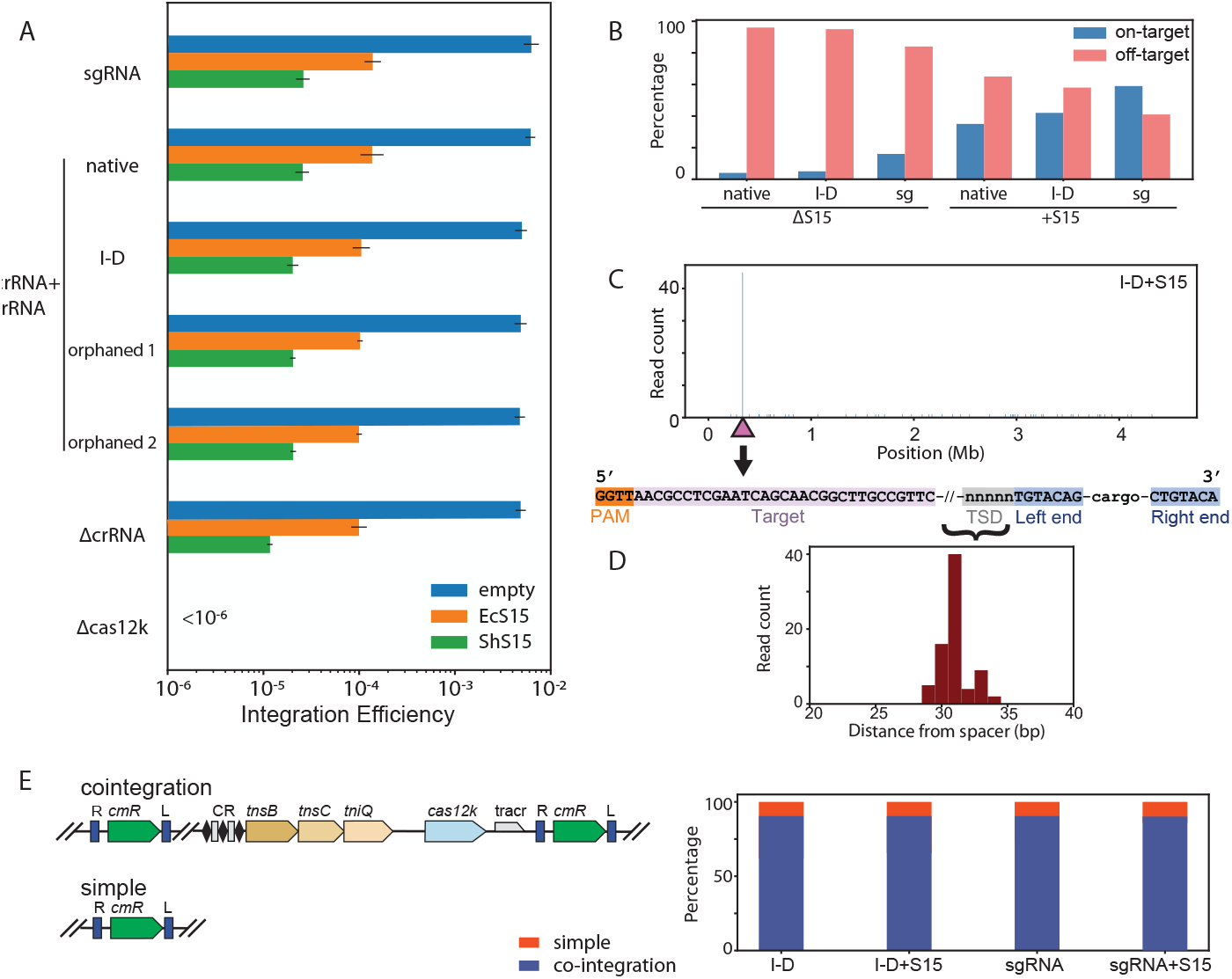
Dual methods of horizontal transmission by Type V CASTs. (A) The small ribosomal protein S15 reduces overall transposition. (B) Over-expression of *E. coli* S15 stimulates on-target transposition, even with a defense-associated type I-D crRNA and the sgRNA. However, significant off-target transposition remains even when S15 is over-expressed. (C) Long-read sequencing with the type I-D crRNA confirms that most, but not all, insertions are in *lacZ* when ShS15 is over-expressed. Triangle: target site. The target site duplication (TSD, gray) and left and right inverted repeats (blue) are also visible at the insertion site. (bottom) (D) The cargo is inserted ∼ 35 − 42 bp away from the target site (purple in B). (E) Schematic (left) and quantification (right) of simple insertion and co-integration products via long-read sequencing. Over-expressing S15 and using heterologous crRNA does not significantly alter this ratio.

## Discussion

Here, we show that CASTs can co-opt defense associated CRISPR arrays already present in the host (Figure 6). This allows CASTs to use the spacers in active defense systems that continuously update their own CRISPR arrays with a record of prior infections. The most recent mobile genetic elements are inserted proximal to the leader of the CRISPR array and are expressed at the highest levels, providing an ample source of crRNAs for the CAST to use for horizontal gene transfer [28]. Type V CASTs also integrate non-specifically via a crRNA-independent mechanism that requires *cas12k*. Type V CASTs are exclusively found in cyanobacteria, suggesting limited horizontal transmission [4, 15]. This may be due to the dependence on S15 for on-target integration, the limited range of the homing spacer, and the possible host toxicity associated with random integration. Analogously to Tn7, type V CASTs may also retain a *tnsE* -like mechanism to target replicating MGEs [6, 8–11]. This work also highlights the need to minimize random integration by type V systems for gene editing applications.

**Figure 6:**
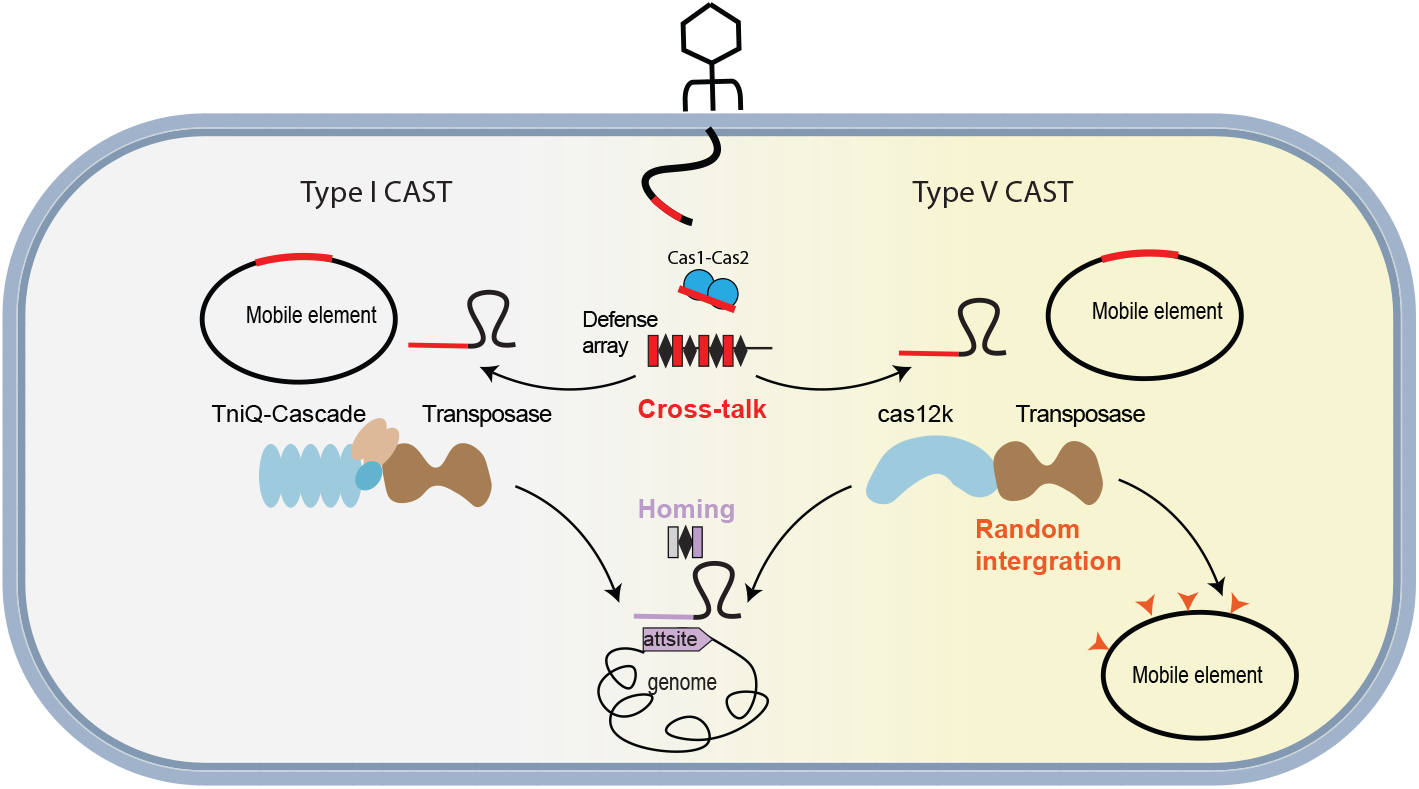
Dual strategies for horizontal transmission by type I and type V CASTs. All CASTs co-opt defense associated CRISPR arrays (red) for horizontal transmission. These arrays are updated by defense-associated Cas1-Cas2 integrases. Type V CASTs can also integrate non-specifically via crRNA-independent transposition. Both systems use a homing spacer for vertical transmission (purple).

Our bioinformatic analysis shows that all type I-B systems co-exist with at least one active CRISPR defense system that can be co-opted by the CAST for horizontal gene transfer. However, we only detected additional CRISPR arrays in ∼ 15% of genomes that harbor a type I-F CAST. Several possibilities may explain this observation. First, type I-F CASTs were primarily found in short genomic assemblies, indicating that these are not high-quality closed genomes. Our analysis may also be oversampling type I-F systems from a small group of cultivatable organisms that are over-represented in the NCBI database. These considerations can bias the results towards fewer systems with co-occurring CRISPR arrays. Second, CASTs may recognize invading DNA by interacting with replisome associated DNA structures (e.g., the replication fork) or other replisome components (e.g., the sliding clamp). For example, Tn7 encodes *tnsE*, a gene that directs transposition to newly-replicated DNA [6, 51]. Third, plasmids and phages also encode their own CRISPR arrays and even full CRISPR systems [52]. These CRISPR arrays can be co-opted by CASTs for mobilization. Additional genome-resolved metagenomic studies, as well as a renewed focus on interactions between CASTs and host proteins, will shed light on these hypotheses.

A type I-F CAST reconstituted with type I-E, I-F, and III-B crRNAs mobilizes via on-target transposition with high integration efficiency. In addition, these CASTs also recognize a wide range of PAMs [53]. Indeed, type I-F CASTs recognize a broader range of PAMs than defense-associated type I-F systems [54]. Type I-F Cascades can also tolerate variable crRNA lengths by adjusting the number of Cas7 repeats in the complex [55–59]. We speculate that type I-F CASTs may also assemble for a variable number of Cas7 subunits, especially on heterologous crRNAs. Taken together, the extraordinary plasticity of type I CASTs in using various spacers and recognizing diverse PAMs facilitates horizontal transmission using defense-associated CRISPR arrays. Conversely, the non-canonical CAST I-F homing spacer prevents defense-associated systems from targeting the host’s genome.

Defense-associated CRISPR sub-types can also share crRNAs. For example, type III systems lack *cas1* and *cas2* and co-occur in genomes containing type I CRISPR-Cas loci [60]. Type III systems can use the pre-processed crRNAs from type I-F systems, acting as secondary defenses that counteract viral escape [61, 62]. Another example is the type VI-B system of *Flavobacterium columnare*, which is also acquisition-deficient. This system can acquire spacers in *trans* from a type II-C system that is encoded in the same genome [63]. CASTs may also use heterologous *cas1* and *cas2* for spacer acquisition into their own arrays. However, CAST CRISPR arrays are extremely short, suggesting that their expansion is not a major mechanism for horizontal transmission. The plasticity of spacer acquisition between CRISPR sub-types suggests that other *cas1/cas2* -deficient systems may use similar mechanisms to target viral pathogens.

Type V CASTs can randomly transpose via a mechanism that does not require a CRISPR array but is dependent on Cas12k. The small ribosomal protein S15 suppresses, but does not completely abrogate, random transposition. Mechanistically, S15 forms a complex with Cas12k and TniQ to stabilize the R-loop [49, 50]. In agreement with earlier work, S15 also increases on-target insertion in the mate-out transposition assay [49, 50]. Interestingly, over-expressing S15 suppressed overall integration via an unknown mechanism. Our assays are conducted *in vivo* and target the *E. coli* genome, which provides ample off-target sites and structures. Additional host factors may further participate in transposition *in vivo* as compared to the purified *in vitro* system [49]. Preventing this off-target transposition activity while boosting on-target integration will be increasingly important for domesticating type V CASTs in heterologous organisms.

## Materials and Methods

### Bioinformatic analysis of CAST co-occurrence with other CRISPR **systems**

Genomes containing CAST systems were collected from NCBI genomic databases [29]. We searched for CRISPR-Cas systems in these genomes using Opfi, a Python library to search DNA sequencing data for putative CRISPR systems [64]. First, we located all regions containing a CRISPR array that was not associated with a CAST. Within those regions, we next searched for *cas* genes located no more than 25 kilobase pairs away from CRISPR array using BLAST and a previously-developed database of diverse *cas* genes [20, 30]. We sub-typed CRISPR-cas systems based on signature genes [60, 65].

### Proteins and nucleic acids

Oligonucleotides were purchased from IDT. Gene blocks for CRISPR arrays were purchased from Twist Biosciences. The R6K plasmid for mate-out transposition assays was obtained from Addgene (#64968) [66]. The type I-F *Vibrio cholerae* HE-45 CAST was sub-cloned from Addgene (#130637 and #130633) [14]. The type V *Scytonema hofmanni* (Sh)CAST was obtained from Addgene (#127922) [15]. The type I-B *Anabaena variabilis* ATCC 29413 CAST was sub-cloned from Addgene (#168137) [16]. For mate-out transposition assays, each of these systems was PCR amplified and cloned into pTNS2 to replace the parental mini-Tn7 (Addgene #64968) by Golden Gate assembly. The repeat, spacer, chloramphenicol resistance cargo, and left and right inverted repeats were synthesized by IDT and cloned into the same plasmid. Full plasmids information can be found in Table s1.

### Cascade purification

Plasmids for type I-F CASTs Cascade over-expression were constructed by sub-cloning the individual genes into pRSFDuet1 (Addgene #126878) to create pIF1008. Type I-F Cascade was co-expressed with 6xHis-MBP-TEV-TniQ and a type III-B crRNA in NiCo21 cells (NEB). Cells were then induced with 0.5 mM isopropyl at 18 ^*°*^C for another 18–20 hours before harvesting. Cells were centrifuged and re-solubilized in lysis buffer containing 25 mM Tris pH 7.5, 200 mM NaCl, 5% glycerol, and 1 mM DTT. Cascade complexes were purified via the N-terminal maltose binding protein (MBP) tag using amylose beads (NEB) and eluted with lysis buffer containing 10 mM maltose. MBP was removed using TEV protease at 4 ^*°*^C overnight. The sample was further diluted to 100 mM NaCl and developed over an anion exchange column (5 mL Q column HP). After loading Cascade, the column was washed extensively with buffer A (25 mM Tris pH 7.5, 100 mM NaCl, 5% glycerol, and 1 mM DTT). The complex was eluted with a 25 column volume gradient of buffer B (25mM Tris pH 7.5, 1 M NaCl, 5% glycerol, and 1 mM DTT.) Cascade was further purified by size exclusion chromatography using a Superose 6 increase column (GE healthcare) in SEC buffer (25 mM Tris pH 7.5, 200 mM NaCl, 5% glycerol, and 1 mM DTT). Fractions were further pooled and concentrated to 0.25 mg/ml and stored in the *−*80 ^*°*^C freezer.

### Cryo-electron microscopy (cryo-EM)

#### Sample preparation and data collection

Purified TniQ-Cascade complexes was diluted to a concentration of 0.25 mg/ml in 25 mM Tris pH 7.5, 200 mM NaCl, 5% glycerol, and 1 mM DTT. Samples were deposited on an Ultra Au foil R 1.2/1.3 grid (Quantifoil) that was plasma-cleaned for 1.5 min (Gatan Solarus 950). Excess liquid was blotted away for 4 s in a Vitrobot Mark IV (FEI) operating at 4 ^*°*^C and 100% humidity before being plunge-frozen into liquid ethane. Data were collected on a Glacios cryo-transmission electron microscope (TEM; Thermo Fisher Scientific) operating at 200 kV, equipped with a Falcon IV direct electron detector camera (Thermo Fisher Scientific). Movies were collected using SerialEM at a pixel size of 0.94Å with a total exposure dose of 40*e*^−^/Å^2^.

#### Data processing and model building

Motion correction, contrast transfer function (CTF) estimation, and particle picking were all performed on cryo SPARC live and further transferred to cryoSPARC for two-dimensional (2D) classification, ab initio 3D reconstruction calculation, 3D classification, and nonuniform refinement [67]. Because of the flexibility of TniQ and Cas6, particle subtraction and focused refinement were also performed in cryoSPARC. A full description of the cryo-EM data processing workflows can be found in Figure S3. A published Cascade structure (PDB: 6PIG) was docked into cryo-EM density maps using Chimera before being refined in Coot, ISOLDE, and PHENIX [46, 68–70]. Full cryo-EM data collection and refinement statistics can be found in Table S2.

#### Conjugation-based transposition assays

CASTs were cloned into a conditionally replicative R6k plasmid (Addgene #64968). The CAST I-F system’s proteins, CRISPR array, and inverted repeats were sub-cloned from Addgene plasmids #130637, #130634, and #130633 to generate pf1001. CAST V and inverted repeat constructs were sub-cloned from Addgene plasmids #127922 and #127924 to generate pIF1005. CAST I-B system’s proteins and inverted repeat constructs were sub-cloned from Addgene plasmids #168137 and #168146 to generate pIF1003.

For transposition, the R6k plasmid was transformed into MFDpir cells, which contain the genomically-integrated RP4-based transfer machinery, termed the donor strain [39]. All growth steps were conducted at 37 ^*°*^C. The donor strain was grown with 0.3 mM diaminopimelic acid (DAP) and appropriate antibiotics. The recipient strain was grown in lysogeny broth (LB). The donor and recipient cells were gently washed four times in PBS (137 mM NaCl, 2.7 mM KCl, 10 mM Na_2_HPO_4_, 1.8 mM KH_2_PO_4_) by spinning and re-suspending 1 mL cultures. The cell density was estimated by taking an optical density reading after re-suspension and the donor and recipient cells were combined in a 3:1 ratio. This mixture was plated on a non-selective plate containing DAP (0.3 mM) for conjugation. The conjugation plate was incubated overnight. The conjugation mixture was collected and washed by mixing with 1 mL PBS, vortexed, and gently spun down four times. Multiple ten-fold dilutions of this mixture were plated onto selective (LB+;12 *μ*l/ml chloramphenicol) and non-selective plates. The cfu/ml was calculated by counting the colonies on plates with 50-500 colonies. The integration efficiency equal to the cfu/ml on selection plate divide by cfu/ml on non-selection plates.

### DNA sequencing of transposition products

#### Sanger sequencing

Individual colonies were re-suspended in LB, pelleted by centrifugation at 16,000 g for 1 min and re-suspended in 80 *μ*l of H_2_O, before being lysed by incubating at 98 ^*°*^C for 10 min in a thermal cycler. Cell debris was pelleted by centrifugation at 16,000 g for 1 min, and the supernatant was removed and serially diluted with 90 *μ*l of H_2_O to generate lysate dilutions for PCR analysis. PCR products were generated with Q5 Hot Start High-Fidelity DNA Polymerase (NEB) using 1 *μl* of the diluted lysate per 10 *μ*l reaction volume. Reactions contained 200 *μ*M dNTPs and 0.5 *μ*M primers and were subjected to 30 thermal cycles. PCR amplicons were resolved by 1% agarose gel electrophoresis and visualized by staining with ethidium bromide (Thermo Scientific). To map integration sites by Sanger sequencing, bands were excised after separation by gel electrophoresis, DNA was gel-extracted (Qiagen), and samples were submitted to Sanger sequencing (Eton).

#### High-throughput long-read whole genome sequencing

Colonies from plate-based transposition reactions were washed off and diluted to an *OD*_600_ ∼ 0.5 using LB. The liquid culture was then grown for two hours at 37 ^*°*^C. Genomic DNA was extracted (ProMega Wizard Genomic DNA kit) and barcoded with a Nanopore technologies rapid barcoding kit. The barcoded DNA was sequenced on a MinION nanopore sequencer using the manufacturer-suggested protocol. Output reads were analyzed using *seqkit* [71]. First, the reads were processed by *seqkit* to collect adjacent target-DNA sequences from all the reads containing 40 bp of the shCAST left end sequence. Then these sequences were mapped onto the BL21(DE3) genome using BLAST+with a 95% sequence identity cutoff [30]. The position of the transposition reaction was defined as the number of basepairs between the end of the PAM and the beginning of the shCAST left-end sequence.

#### Single-colony whole genome sequencing

Single colonies were grown overnight in LB with 17 ng/*μ*l chloramphenicol at 37 ^*°*^C. 1 ml of the overnight culture was pelleted by centrifugation at 16,000 g for 1 min and the gDNA was extracted asb described above. Genomic DNA samples were separated and barcoded in units of 12 per batch using the MinION rapid barcoding kit. Samples were loaded into a MinION flowcell (FLO-MIN106D) and sequenced with a MinION Mk1B device. The raw read fastq files were assembled with flye [72].

## Supplemental Information

Supplemental information includes seven figures and two tables.

## Declarations

## Author Contributions

K.H. and I.J.F. conceived the project. K.H., C.-W.C., and Z.Y. performed all experiments and analyzed the data. K.H. and C.-W.C. prepared the figures. I.J.F. and C.O.W. secured the funding. I.J.F. and C.O.W. supervised the project. K.H., C.-W.C., C.O.W. and I.J.F. wrote the manuscript with input from all co-authors.

## Funding

This work was supported by NIGMS grants R01GM124141 (to I.J.F.) and R01GM088344 (to C.O.W.), the Welch Foundation grant F-1808 (to I.J.F.), and the College of Natural Sciences Catalyst Award for seed funding.

## Declaration of Interests

K.H., C.O.W. and I.J.F. have filed a patent application relating to CRISPR-associated transposons.

## Notes

### Competing Interest Statement

The authors have declared no competing interest.

### Summary of Updates

With the adjustment of tracrRNA expression, we showing the type V CAST systems also doing cross-talking like type I-F CAST systems.

