## Supplemental Table 1-2, Supplemental Figure 1-6 for "Distinct horizontal transfer mechanisms for type I and type V CRISPR-associated transposons"

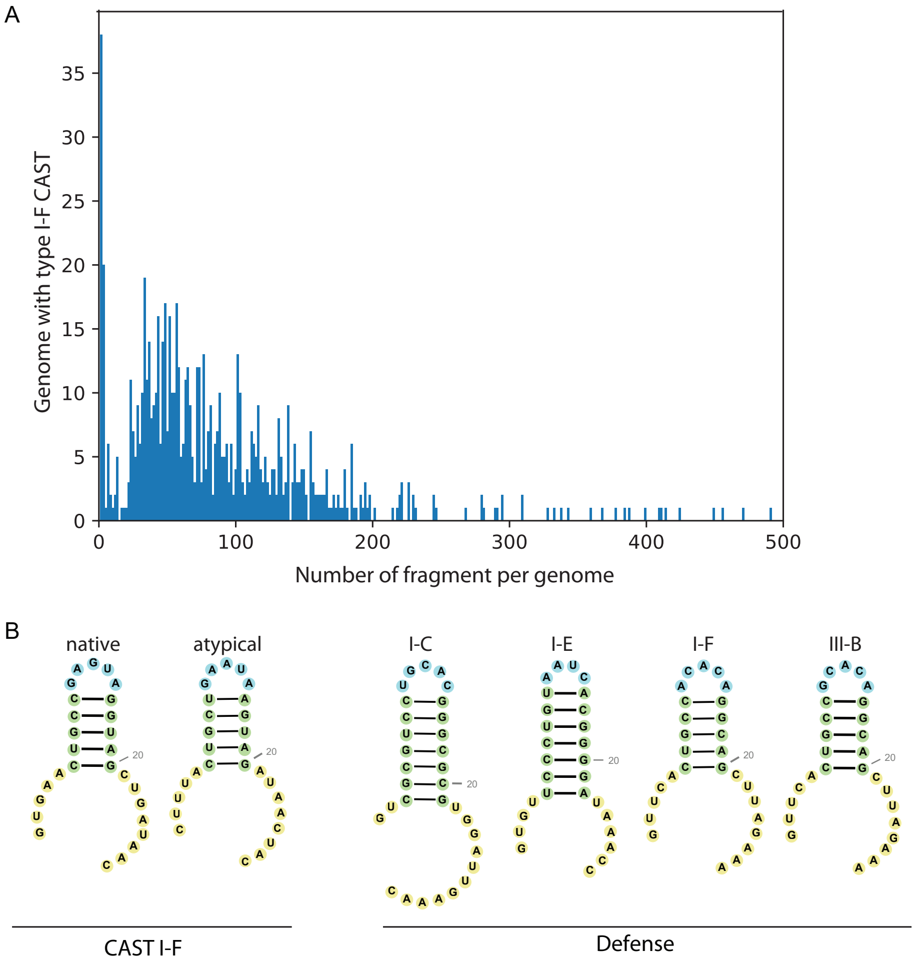

C

| | native | atypical | Defense<br>I-C | I-E | I-F | III-B |
|---|---|---|---|---|---|---|
| blue | 0 | 0 | N.A. | 2 | 0 | 0 |
| white | 395 | 181 | N.A. | 43 | 167 | 401 |

**Figure S1: A type I-F CAST catalyzes on-target transposition using heterologous CRISPR arrays.** (A) The large number of contigs in genomes that encode CAST I-F systems indicates that these may not be complete assemblies. This suggests that the number of co-occurring CAST and defense systems is a lower bound on the true co-occurrence frequency. (B) The structure of native type I-F CAST direct repeats (DRs) compared to the structures of DRs from co-occurring CRISPR-Cas systems. Colors indicate different structural elements in the DR. Blue: loop; green: stem; yellow: handles. (C) On-target transposition into *lacZ* results in white colonies on LB+CmR+X-gal plates. In this example, the type I-F CAST and its native crRNA are guided to *lacZ* in the recipient cell genome. Integration is confirmed via junction PCR and Sanger sequencing.

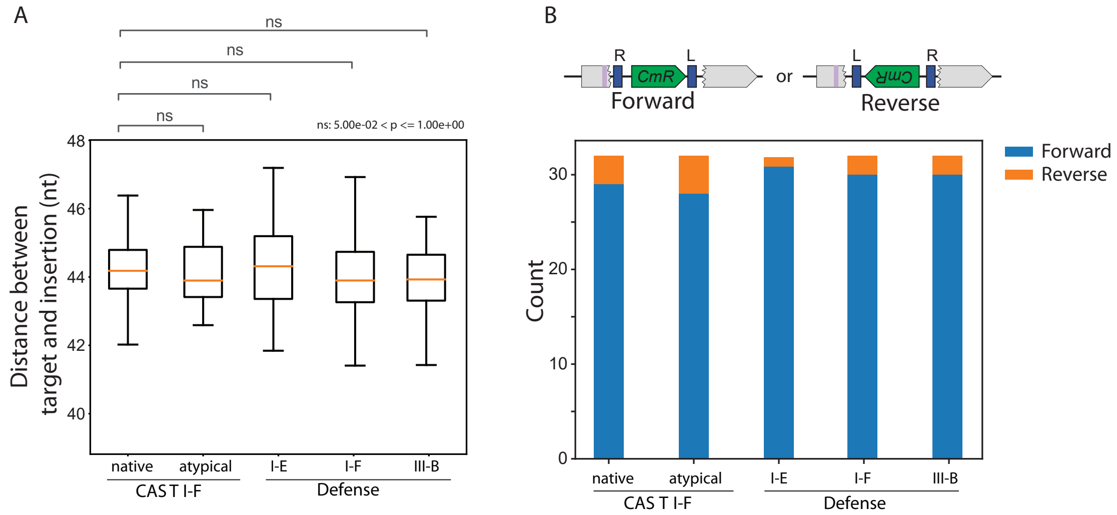

**Figure S2: Defense-associated CRISPR RNAs support on-target integration.** (A) The distance between the target site and the integration site for the indicated crRNAs, as determined via Sanger sequencing of 32 colonies for each condition. n.s.: $p > 0.05$. (B) Schematic of forward and reverse integration orientations (top). The integration orientation was determined via Sanger sequencing for each of the indicated crRNAs (bottom).

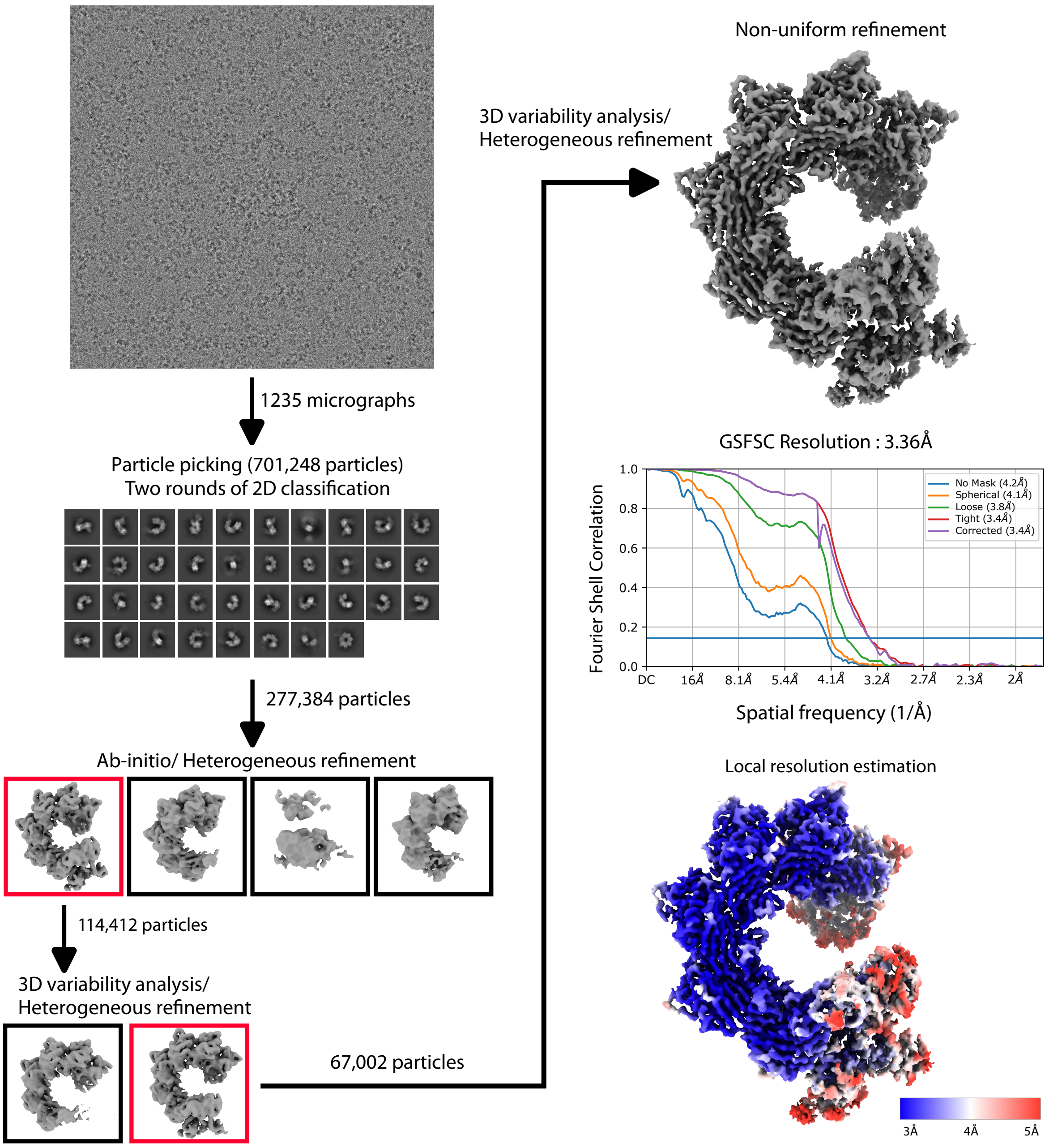

**Figure S3: CryoEM image processing workflow for the TniQ-Cascade with a defense-associated type III-B crRNA.**

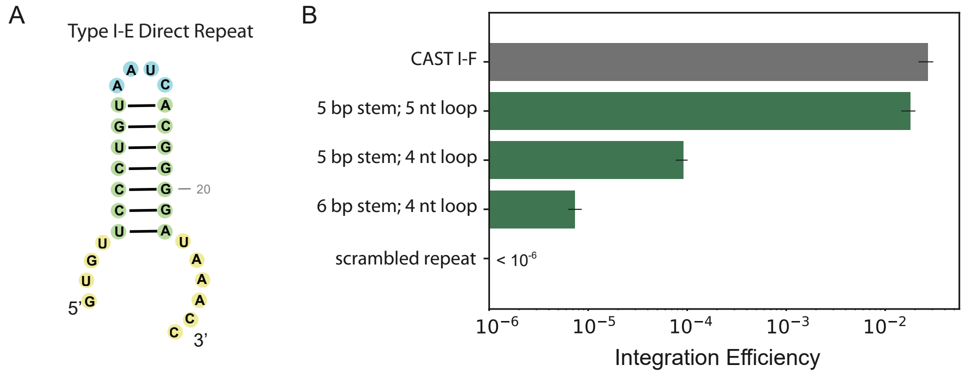

**Figure S4: The length of the direct repeat (DR) stem loop controls transposition.** (A) Predicted structure of the DR from a defense-associated type I-E CRISPR-Cas system. (B) Changing the loop and stem length increases integration efficiency. Error bars: mean of three replicates.

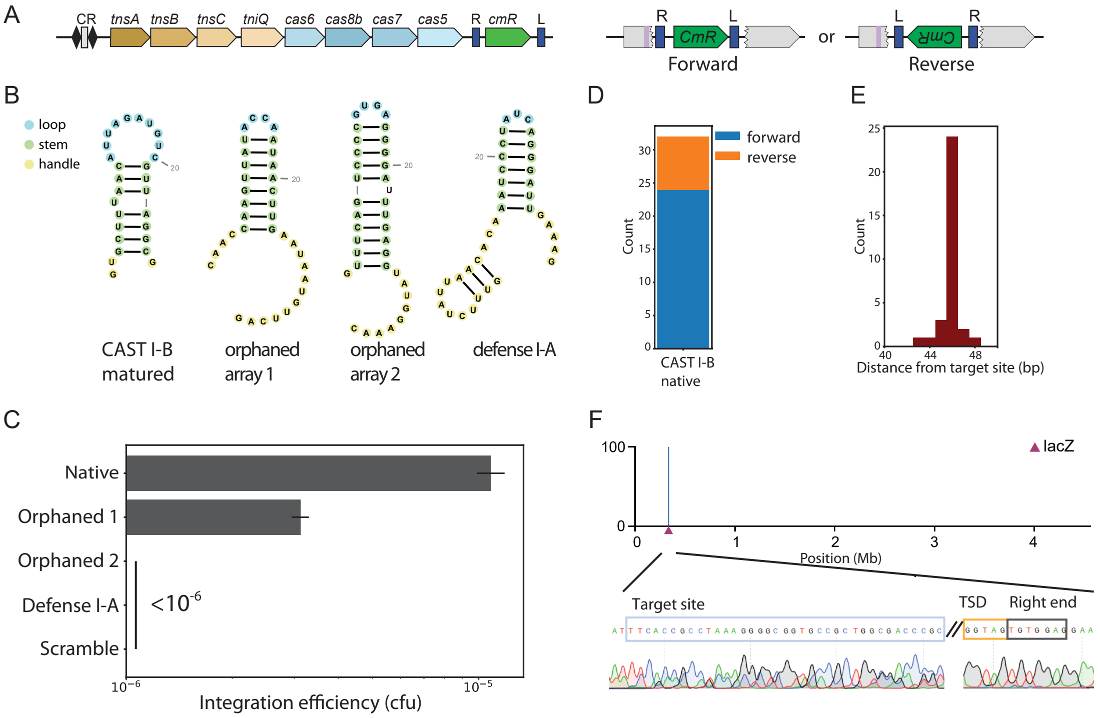

**Figure S5: Type I-B CASTs co-opt spacers from heterologous CRISPR arrays.** (A) Gene architecture of the type I-B *Av*CAST, cloned into the R6k plasmid (pIF1004, Table S1). (B) Predicted structure of the direct repeat (DR) from all CRISPR arrays found in *Anabaena variabilis* (ATCC 29413). (C) Quantification of transposition from the native CAST array and co-occurring CRISPR systems. Error bars: standard deviation across three biological replicates. Scrambling the DR suppressed transposition below our detection limit of $\sim 10^6$ cfus. (D) The integration orientation and (E) distance between the target and insertion sites were determined via Sanger sequencing (N=32 clones). (F) Long-read NGS confirms single on-target cut-and-paste transposition into *lacZ*. Target site duplication (TSD) is also visible in this data.

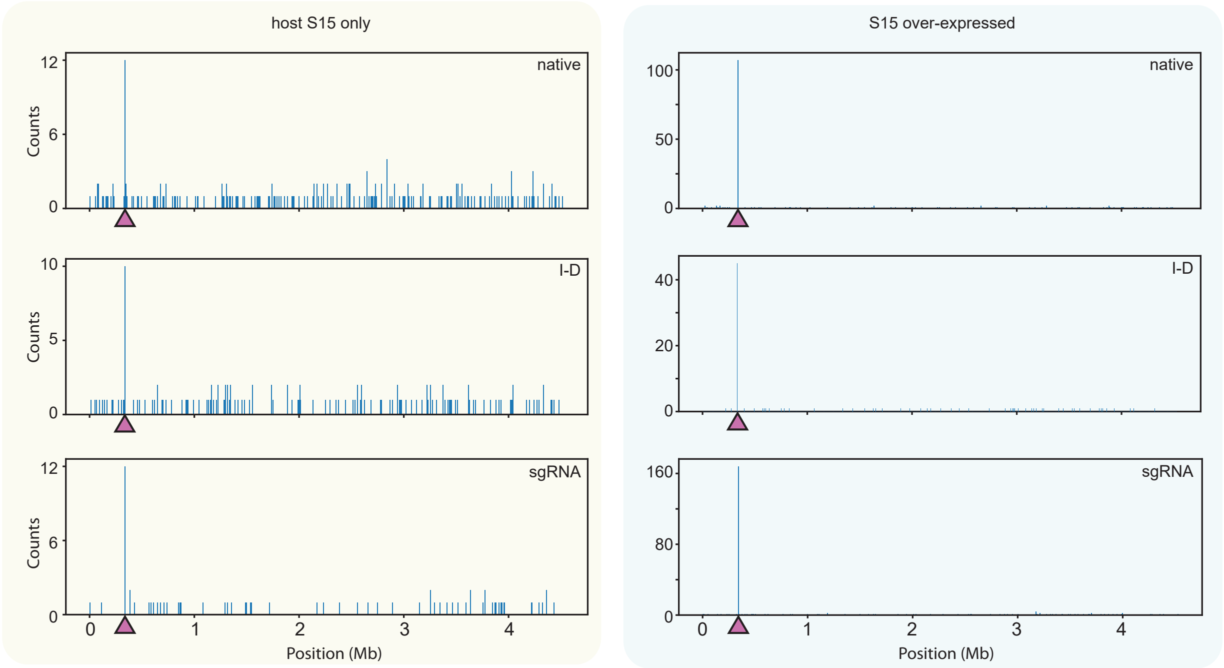

**Figure S6: Long-read sequencing of transposition events by a type-V CAST with a native crRNA (top), type I-D crRNA (middle), or a sgRNA (bottom).** Left: S15 is not over-expressed; Right: S15 is over-expressed in the recipient cells. Triangle: target site.

### Supplemental Tables

| Plasmid | Description | Source |
|---|---|---|
| pIF1010 | miniTn7-encoding R6K plasmid | Addgene: 64968 [1] |
| pIF1011 | *V. cholerae* TniQ, Cas8, Cas7, and Cas6 | Addgene: 130637 [2] |
| pIF1012 | Kanamycin resistance cassette | Addgene: 130634 [2] |
| pIF1013 | *V. cholerae* TnsA, TnsB, and TnsC | Addgene: 130633 [2] |
| pIF1014 | ShCAST system and its native crRNA | Addgene: 127922 [3] |
| pIF1015 | Kanamycin resistance cassette for ShCAST | Addgene: 127924 [3] |
| pIF1016 | Expression of AvCAST proteins | Addgene: 168137 [4] |
| pIF1017 | AvCAST donor | Addgene: 168145 [4] |
| pIF1008 | R6K backbone with Golden Gate (GG) cloning sites | This paper |
| pIF1001 | R6K plasmid encoding a type I-F CAST but no CRISPR array | This paper |
| pIF1002 | R6K plasmid with a I-F CAST targeting *lacZ* | This paper |
| pIF1003 | R6K plasmid with a I-B CAST but no CRISPR array | This paper |
| pIF1004 | R6K plasmid with a I-B CAST targeting *lacZ* | This paper |
| pIF1005 | R6K plasmid encoding a type V CAST but no CRISPR array | This paper |
| pIF1006 | R6K plasmid with a type V CAST targeting *lacZ* | This paper |
| pIF1008 | Type I-F Cascade over-expression | This paper |
| pIF1009 | Expression of a type III-B defense-associated crRNA | This paper |

**Table 1:** Plasmids used in this study.

| **Data collection and Processing** | |
|---|---|
| Microscope | Glacios |
| Pixel size (Å) | 0.94 |
| Voltage (kV) | 200 |
| Detector | Falcon IV |
| Exposure ($e^{-}/\text{Å}^2$) | 40 |
| Defocus range ($\mu$m) | -1.2 to -2.2 |
| Final particles | 66,862 |
| Symmetry | C1 |
| Map Resolution (Å) (FSC threshold=0.143) | 3.36 |
| **Model refinement and validation** | |
| Initial model used | 6PIG |
| Ramachandran | |
| Flavor (%) | 95.63 |
| Allowed (%) | 4.37 |
| Outlier (%) | 0.00 |
| Rotamer outliers (%) | 0.16 |
| Clash score | 7.88 |
| MolProbity score | 1.73 |
| EM Ringer score | 2.82 |

**Table 2:** Cryo-EM data collection and processing statistics
